# Deformed wing virus genotypes A and B do not elicit immunologically different responses in naïve honey bee hosts

**DOI:** 10.1101/2023.08.26.554974

**Authors:** Amanda M Norton, Gabriele Buchmann, Alyson Ashe, Owen T Watson, Madeleine Beekman, Emily J Remnant

**Author notes:** Present address: Research and Advanced Instrumentation, Academic Support Unit, University of the Sunshine Coast, Sippy Downs, Queensland, Australia (Amanda M Norton). Present address: Institute of Plant Genetics, Heinrich-Heine University, Duesseldorf, Germany (Gabriele Buchmann). Address correspondence to Emily J. Remnant.

## Abstract

*Deformed wing virus* (DWV), in association with *Varroa destructor*, is currently the leading factor associated with global honey bee deaths. With the exception of Australia, the virus and mite have a near global distribution, making it difficult to separate the effect of one from the other. Over time, the prevalence of the two main DWV genotypes (DWV-A and DWV-B) has changed, leading to the suggestion that the two strains elicit a different immune response by the host, the western honey bee *Apis mellifera*. Here we use a honey bee population naïve to both the mite and the virus to investigate if honey bees show a different immunological response to DWV genotypes. We examined the expression of 19 immune genes by RT-qPCR and comprehensively analysed the small RNA response in honey bees after experimental injection with DWV-A and DWV-B. We found no evidence to indicate that DWV-A and DWV-B elicit a different immune response in honey bees. We found that RNA interference genes are up-regulated during DWV infection and that the small interfering RNA (siRNA) response is proportional to viral loads, yet does not inhibit the virus from accumulating to high loads. We also found that the siRNA response towards DWV was weaker than the response to another honey bee pathogen, *Black queen cell virus*. This suggests that DWV is comparatively better at evading antiviral host defences. There was no evidence for the production of virus-derived PIWI-RNAs in response to DWV infection. In contrast to previous studies, and in the absence of *V*. *destructor*, we found no evidence that DWV has an immunosuppressive effect in honey bees. Overall, our results advance our understanding of the immunological effect DWV elicits in honey bees.

## 1. INTRODUCTION

Obligate pathogens need to evade or suppress host immune defences in order to hijack the host’s cellular machinery. The success of the pathogen largely depends on how effectively it can enter and replicate within a host cell, and conversely, how successfully the host can inhibit, degrade or tolerate the infectious agent (Hedrick, 2017). Avoidance is often the first defence strategy against an infectious agent (Curtis, 2014; Townsend et al., 2020). Disease avoidance through the use of quarantine was first practiced in 14^th^ century, whereby ships carrying sick individuals were prohibited from docking at port in Venice, Italy, for 40 days to stop the transmission of the bubonic plague (Kilwein, 1995). Avoiding transmission can be challenging when living in dense populations with frequent social interactions, due to the increased probability of encountering an infected individual. Rapid human population growth in cities, overcrowded slums, poor sanitation and contaminated water supplies, create ideal conditions for the transmission of human diseases such as tuberculosis, typhoid and cholera (Duffy, 1971).

Social insects such as ants, termites, and some bee and wasp species, which live in often densely populated colonies, are particularly vulnerable to infectious disease. Pathogen avoidance is the first defence against disease transmission. Social insects have numerous behavioural adaptations that collectively provide ‘social immunity’ (Cremer et al., 2018; Simone-Finstrom, 2017), such as corpse removal in bees (Rosengaus et al., 1999) and ants (Wilson et al., 1958), pathogen alarm signalling in termites (Rosengaus et al., 1999), use of plant resins as antimicrobial agents in bees (Simone et al., 2009), and specific worker castes to deal with waste in ants (Hart and Ratnieks, 2001). These behavioural responses are so effective that they are thought to have contributed to the reduction in the number of immune genes that social insects have compared to their solitary counterparts (Harpur and Zayed, 2013). Nonetheless, social insects still rely upon an innate immune response at the individual level, as even the most hygienic colonies cannot entirely avoid pathogens.

Social insects have a number of immune response pathways involved in dealing with pathogens. Small interfering RNAs, operating within the RNA interference (RNAi) pathway, are the major antiviral defence mechanism. Double stranded RNA produced during the replication of viruses is recognised by the protein dicer, which subsequently cleaves the RNA into 21-23 nt fragments termed virus-derived small interfering RNAs (vsiRNAs). The vsiRNAs bind to an argonaute protein and are loaded into an RNA-induced silencing complex (RISC), which catalyses the degradation of targeted viral RNA (Hammond et al., 2001). The Toll, Imd and JAK/STAT pathways are microbial sensing pathways that detect pathogens via cell surface receptors and initiate a downstream transcriptional response (Lemaitre and Hoffmann, 2007). The Toll pathway is generally associated with host response to fungi and Gram-positive bacteria, and the Imd pathway with Gram-negative bacteria. *Dorsal* (Toll), and *relish* (Imd) are both NF-κB family transcription factors, involved in the production of antimicrobial peptides (AMPs) in fat bodies, a tissue analogous to the mammalian liver (Lemaitre and Hoffmann, 2007). The Toll and Imd pathways may also be involved in antiviral defence (Brutscher et al., 2015).

Parasites and pathogens are a particularly serious problem for the Western honey bee *Apis mellifera,* due to a global reliance on honey bees as a pollinator of commercial crops. Colony densities are unnaturally high in many areas, such as almond-growing regions in Australia and the US, where approximately 200,000 and 2,000,000 honey bee colonies are moved annually for the purpose of pollination (Le Feuvre, 2017; Lee et al., 2018). In addition, beekeeping practices lead to increased colony sizes, as bigger colonies are better pollinators and produce more wax and honey (Farrar, 1937). This facilitates the transmission of parasites and pathogens within and between colonies (Alger et al., 2018; Brosi et al., 2017).

The dynamics of one pathogen in particular, *Deformed wing virus* (DWV), were considerably altered when the ectoparasitic mite *Varroa destructor* shifted hosts from the Asian honey bee (*Apis cerana*) to *A. mellifera* (Martin, 2001). *V. destructor* is a competent vector of RNA viruses such as DWV (Allen and Ball, 1996) facilitating increased transmission within and between honey bee colonies. DWV has spread across the globe over the last 40 years, driven by the spread of *V. destructor* (Wilfert et al., 2016). Until June 2022, Australia was the last major beekeeping nation to remain free of *Varroa* (Chapman et al., 2023; Martin and Brettell, 2019; Roberts et al., 2017). While much of the Australian continent remains unaffected, an incursion zone has been established and eradication is being pursued by authorities. The combination of *V. destructor* and DWV is considered to be the predominant driver of global honey bee colony deaths (Dainat et al., 2012; Martin, 2001; Schroeder and Martin, 2012). Australia remains free of DWV (Roberts et al., 2017) and there is no evidence it was introduced in the current *V. destructor* incursion (Chapman et al., 2023).

The majority of Deformed wing virus infections occur due to two related DWV genotypes, DWV-A and DWV-B, with genome sequences that differ by approximately 15% (Ongus et al., 2004). These DWV genotypes also appear to differ in their effect on honey bees, but many studies find contradictory results. Some studies report increased mortality in colonies that harbour high DWV-A loads (Barroso-Arévalo et al., 2019a; Highfield et al., 2009; Kevill et al., 2019; Kevill et al., 2017; Martin et al., 2012; Mondet et al., 2014), while others report reduced survival and cognitive function in bees that were experimentally injected with DWV-B (Benaets et al., 2017; Gisder et al., 2018; McMahon et al., 2016). Our own earlier work found increased mortality in pupae injected with DWV-A (Norton et al., 2020), while others found no difference in survival in pupae injected with either genotype (Dubois et al., 2019; Tehel et al., 2019).

Many factors can potentially contribute to differences in observed mortality between studies. One explanation is that the bees used between studies differ in their immune response to DWV genotypes. Some honey bee populations may be more susceptible to one genotype of DWV, whereas others may be more susceptible to the other genotype. Such differences could be a result of pre-exposure to a particular strain, resulting in immune priming – improved survival if a sublethal dose of pathogen has been previously encountered (Hernández López et al., 2014). Secondly, it is possible that honey bees elicit a specific and different immune response to different genotypes of the same pathogen. Specific genotype-host immune interactions are known from parasites in bumblebees (Barribeau and Schmid-Hempel, 2013) and mosquitos (Molina-Cruz et al., 2012), bacteria in crustaceans (Auld et al., 2012) and humans (Sela et al., 2018), and viruses in fish (Moreno et al., 2020).

Here we ask if honey bees mount a different immune response when experimentally injected with DWV-A compared to DWV-B. To test for differing responses, we used a honey bee population naïve to DWV and *V. destructor*. Our study thus minimises confounding factors such as exposure or adaptation to a particular DWV genotype, or direct damage inflicted by mite feeding (Annoscia et al., 2019; Kuster et al., 2014; Ramsey et al., 2019; Yang and Cox-Foster, 2005). We analysed the expression of 19 genes that represent several immune response pathways and have previously been associated with DWV (Table 1). We additionally sequenced the small RNA profiles of honey bees infected with DWV-A and DWV-B to better understand the role of RNAi during DWV infection.

**Table 1.** Immune gene expression in DWV infected honey bees in this study compared to previous studies. Treatment indicates if bees were parasitised by *V*. *destructor* or collected from mite-infested colonies (DWV+), or experimentally injected (DWV*i*) in past studies. Asterisks indicate variability in expression across time points, and in our study this is in comparison to the buffer injected or unmanipulated controls.

| Pathway | Gene | Past studies |  |  | This study | References |
| --- | --- | --- | --- | --- | --- | --- |
|  |  | Treatment | life stage | expression | expression |  |
| RNAi | *dicer* | DWV+ | adults | up-regulated | up-regulated | (Zhao et al., 2019)* |
|  |  | DWVi | adults | up-regulated |  | (Al Naggar and Paxton, 2021) |
|  |  | DWV+ | pupae | equal to controls |  | (Ryabov et al., 2014) |
|  | *argonaute-2* | DWV+ | adults | up-regulated | up-regulated* | (Zhao et al., 2019)* |
|  |  | DWVi | adults | up-regulated |  | (Al Naggar and Paxton, 2021) |
|  |  | DWV+ | pupae | equal to controls |  | (Ryabov et al., 2014) |
| Toll | *dorsal-1a* | DWV+ | adults | down-regulated | equal to both controls | (Barroso-Arévalo et al., 2019b; Nazzi et al., 2012)* |
|  |  | DWV+ | pupae | down-regulated |  | (Annoscia et al., 2019) |
|  | *cactus* | DWV+ | adults | up-regulated | mostly equal to controls* | (Zanni et al., 2017) |
|  | *spaetzle* | DWV+ | pupae | down-regulated | equal to buffer inj. | (Khongphinitbunjong et al., 2015) |
|  |  | DWV+ | pupae |  |  | (Ryabov et al., 2014) |
|  | *PGRP-S2* | DWV+ | adults | up-regulated | mostly equal to controls* | (Nazzi et al., 2012) |
|  |  | DWV+ | adults | down-regulated |  | (Zanni et al., 2017) |
| Imd | *relish* | DWV+ | adults | up-regulated | up-regulated* | (Barroso-Arévalo et al., 2019b)* |
|  |  | DWV+ | larvae + pupae | up-regulated |  | (Kuster et al., 2014)* |
|  |  | DWV+ | adults | up-regulated |  | (Barroso-Arévalo et al., 2019b)* |
|  |  | DWV+ | pupae | down-regulated |  | (Khongphinitbunjong et al., 2015) |
| AMPs | *defensin-1* | DWVi | pupae | up-regulated | mostly equal to buffer inj.* | (Ryabov et al., 2016) |
|  |  | DWV+ | pupae |  |  | (Khongphinitbunjong et al., 2015) |
|  |  | DWV+ | adults + pupae |  |  | (Aronstein et al., 2012) |
|  |  | DWV+ | adults |  |  | (Zhao et al., 2019)* |
|  |  | DWV+ | adults | down-regulated |  | (Barroso-Arévalo et al., 2019b)* |
|  | *defensin-2* | DWV+ | larvae + pupae | up-regulated | mostly equal to buffer inj.* | (Kuster et al., 2014)* |
|  | *abaecin* | DWV+ | adults | up-regulated | mostly equal to buffer inj.* | (Zanni et al., 2017; Zhao et al., 2019)* |
|  |  | DWV+ | adults + pupae | no significant change |  | (Aronstein et al., 2012) |
|  | *hymenoptaecin* | DWV *i* | pupae | up-regulated | equal to buffer inj. up to 48 HPI* | (Ryabov et al., 2016) |
|  |  | DWV+ | adults |  |  | (Zanni et al., 2017; Zhao et al., 2019)* |
|  |  | DWV+ | larvae + pupae |  |  | (Kuster et al., 2014)* |
|  |  | DWV+ | adults + pupae | no significant change |  | (Aronstein et al., 2012) |
|  |  | DWV+ | adults | down-regulated |  | (Yang and Cox-Foster, 2005) |
|  | *apidaecin* | DWV+ | adults | up-regulated | mostly equal to buffer inj.* | (Zanni et al., 2017; Zhao et al., 2019) |
|  |  | DWV+ | larvae | down-regulated |  | (Di Prisco et al., 2016) |
| **JAK/STAT** | *domeless* | DWV+ | pupae | down-regulated | up-regulated in DWV-B* | (Khongphinitbunjong et al., 2015) |
|  | *hopscotch* | DWV+ | pupae | down-regulated | up-regulated from 72 HPI * | (Ryabov et al., 2014) |
|  |  | DWV+ | adults | up-regulated |  | (Zhao et al., 2019)* |
| **Other** | *lysozyme* | DWV+ | adults | down-regulated | mostly equal to controls* | (Yang and Cox-Foster, 2005) |
|  | *malvolio* | DWV+ | adults | up-regulated | up-regulated | (Zanni et al., 2017) |
|  | *vago* | DWV+ | pupae | up-regulated | mostly equal to controls* | (Ryabov et al., 2014) |
|  | *PPOact* | DWV+ | pupae | down-regulated | mostly equal to controls* | (Khongphinitbunjong et al., 2015) |

## 2. MATERIALS AND METHODS

### 2.1 Sample preparation

Infectious viral particles used to prepare DWV inocula were obtained from adult bees collected in Blenheim, New Zealand in 2016 (DWV-A; GenBank accession MN538208) and Amsterdam Water Dunes, the Netherlands in 2015 (DWV-B; GenBank accession MN538209). Details of inocula preparation, standardisation and sequencing are given in Norton et al, (2020). We collected white-eyed pupae from three unrelated *A*. *mellifera* colonies in Sydney, Australia, which are naïve to both *V*. *destructor* and DWV. Pupae were assigned to four treatment groups of 95 pupae per colony. Pupae were either left unmanipulated (‘unmanipulated control’; UC), or intra-abdominally injected with a 32G needle and 10 μL Hamilton syringe containing 2 μL 0.5 M potassium phosphate buffer pH 8 (‘buffer control’; BC), or 1 × 10^7^ genome equivalents of DWV-A or DWV-B. All pupae were placed into Petri dishes lined with sterile filter paper (10 pupae/Petri dish) stored in sealed plastic tubs and incubated at 34.5**^°^**C for 8 days (192 hours). During incubation, four random pupae per colony and treatment group were collected at specific time points post-injection (1, 4, 8, 12, and 24 h) and every subsequent 24 h for 192 h. Selected pupae were frozen at −80**^°^**C prior to RNA extraction. RNA extraction of individual pupae and RNA standardisation to 200 μg/mL were conducted as described (Norton et al, 2020). A portion of the RNA was used in our previous study to analyse DWV levels (Norton et al, 2020), and a portion used here to generate cDNA for immune gene analysis. We synthesised cDNA from 800 ng RNA with SuperScript IV VILO Master Mix with ezDNase enzyme (Invitrogen) in 10 μL reaction volumes. The cDNA was then diluted in 150 μL UltraPure nuclease-free dH_2_O prior to qPCR analysis.

Previously, In our previous study, we found only minor differences between colonies in terms of DWV strain accumulation (Norton et al., 2020; see also Figure 5B), so we combined pupae across all three colonies for global immune gene expression analysis. Where possible, we analysed the gene expression of 12 pupae at each time point (4 pupae per treatment / per colony). However, we excluded seven pupae (2.1%) injected with either DWV-A or DWV-B that were contaminated with the other genotype during downstream processing (Norton et al., 2020), and two UC pupae (0.6%) that were contaminated with DWV-A or DWV-B. Pupae were screened for additional viruses by qPCR [*Black queen cell virus* (BQCV), *Sacbrood virus* (SBV), *Apis rhabdovirus* 1 and 2 (ARV-1/ARV-2)] and endpoint PCR (*Lake Sinai virus*) as described (Norton et al., 2020). High loads of BQCV (≥ 1 × 10^7^ relative to *Actin*) were detected in 4.9% of DWV-A injected pupae, despite BQCV not being detected in the sequenced inocula (Norton et al., 2020). Generally, only one pupa per colony/time point had high BQCV loads, excluding 72 HPI, where this occurred in six pupae, and at 96 HPI where three pupae had high BQCV loads. To reduce any confounding effects, we removed pupae contaminated with BQCV or the alternate DWV strain from the immune gene qPCR analyses. Our final treatment groups consisted of 320 total individual pupae: 82 (UC), 84 (BC), 76 (DWV-A) and 78 (DWV-B), so that even with removal of individuals, at least 6-12 individual pupae were analysed per treatment and time point. Our small RNA analysis was conducted by pooling pupae from each colony, rather than individual pupae (below), so we included all four pupae per time point/colony to ensure our pooled samples were the same size.

### 2.2 qPCR analysis

We analysed 19 immune related genes and three endogenous reference genes (*Actin*, *RpL8* and *RpS5*) by qPCR (Table S1) at the 7 different timepoints mentioned above (8, 12, 24, 48, 72, 96 and 192 HPI). All 5 μL qPCR reactions were performed in duplicate in 384-well plates (Bio-Rad) using a Pipetmax 268 (Gilson) with SsoAdvanced Universal SYBR Green supermix (Bio-Rad), forward and reverse primers (final concentration 500 nM each), and 1 μL cDNA. qPCR analysis was conducted on a Bio-Rad CFX384 Touch real-time PCR detection system with cycling conditions 95°C (10 min), followed by 40 cycles of 95°C (15 s), 55-63.2°C depending on primer pair (15 s), and 72°C (10-30 s), and then followed by melt curve analysis between 55 and 95°C, at 0.5°C increments. Each run contained duplicate positive and negative (no template) controls. We validated the specificity of each primer pair with melt curve analysis and gel electrophoresis on 1% agarose gel with SB buffer and SYBR Safe DNA stain (Life Technologies). We calculated primer efficiencies from standard curves prepared from a six-step 10-fold dilution series of cDNA. Primer efficiencies ranged from 95-110%. We determined the stability of the three reference genes with BestKeeper (Pfaffl et al., 2004), and found that *Actin* and *RpS5* were the most stable. Estimates of the expression level of each immune related gene were calculated as *E^CqMin^*−*^Cqi^*and normalised against the geometric mean of the two reference genes (Vandesompele et al., 2002), as described by Brito et al. (2010).

### 2.3 Gene expression statistical analysis

For each given gene, we stratified the expression data by time-point and compared the mean rank of gene expression between the four treatment groups with a Kruskal-Wallis test, followed by Dunn’s posthoc test with Benjamini-Hochberg adjustment [‘FSA’ (Ogle et al., 2018)]. We used non-parametric analysis as parametric assumptions of homogeneity of variance and/or normality were violated for 11 genes, and could not be improved with transformations. All statistical analyses were performed with RStudio software (R version 3.5.0). We generated an expression heatmap with ‘gplots’ (Warnes et al., 2016) to visualise the gene expression patterns across all time points and all genes.

### 2.4 Small RNA library preparation and sequencing

To more closely examine active viral degradation by the immune system via the RNA interference pathway, we sequenced small RNAs in DWV-A and DWV-B injected pupae at 12, 24, 48 and 96 HPI. We chose these specific time points as we were interested in investigating the small RNA response in relation to increasing viral loads. The RNAi mechanism includes three small RNA pathways that may all have a role in antiviral defence: the small interfering (siRNA), PIWI-interacting (piRNA), and microRNA (miRNA) pathways. While the siRNA pathway is the major antiviral response in insects (Gammon and Mello, 2015), virus derived piRNAs are associated with antiviral defence in mosquitos (Goic et al., 2016; Miesen et al., 2016), and host-derived miRNAs can play an antiviral response and be differentially expressed during viral infections (Monsanto-Hearne and Johnson, 2018; Slonchak et al., 2014; Yan et al., 2014). It is unknown if honey bees produce either piRNAs or miRNAs in response to viral infection.

We pooled 0.8 μg RNA from each pupa, to give a single DWV-A or DWV-B sample per colony and time point. We then used 0.6 μg of the pooled RNA to generate small RNA libraries using NEBNext® Multiplex Small RNA Library Prep Set for Illumina. We followed the manufacturer’s protocol with the following modifications: libraries were generated from half-reactions, we used the Qiagen MinElute PCR Purification Kit for PCR purification and adjusted pH with 10 μl 3M sodium acetate pH 5.5. Acrylamide gel separation of the libraries was conducted on a 6% Novex TBE PAGE gel with 5 μL Novex Hi-Density TBE Sample Buffer (5X). Gels were stained with SYBR Gold in TBE buffer for 20 min. The excised gel slices were passed through gel breaker tubes. Gel elution was performed overnight at 4°C. We used 2 µl of glycogen instead of linear acrylamide for precipitation and elution was performed with 10 µl TE elution buffer from the kit. The libraries were sequenced at the Australian Genome Research Facility (AGRF) laboratory (Melbourne, Australia) with HiSeq 2500 100 bp single end sequencing.

### 2.5 Small RNA analysis

#### 2.5.1 siRNA analysis

We checked the quality of the sequencing data with FastQC (http://www.bioinformatics.babraham.ac.uk/projects/fastqc/), and trimmed adapters with Trimmomatic (Bolger et al., 2014). We used consensus DWV-A (accession MN538208), DWV-B (MN538209) reference genomes and a BQCV reference genome (MW390818) assembled *de novo* (see below) to produce index libraries and used Bowtie2 (Langmead and Salzberg, 2012) to align the small RNA reads to either the DWV-A, DWV-B or BQCV genome. The mapped reads were exported as BAM files using SAMtools (Li et al., 2009), and analysed with the R package viRome (version 0.10).

#### 2.5.2 piRNA analysis

We filtered the mapped reads to remove reads less than 26 and greater than 31 nucleotides in length, corresponding to the expected lengths of piRNAs (Aravin et al., 2006). SAMtools was used to remove duplicate reads for analysis of unique piRNA species and for BAM to fastQ file conversion (Li et al., 2009). The nucleotide frequencies for unique and total reads of each sample were plotted with R package SeqTools (Barson and Griffiths, 2016).

#### 2.5.3 miRNA analysis

FastQ files were imported into CLC Genomics and passed through quality control and trimming tools. Differential expression analysis was conducted by performing the extract and count tool, followed by annotate and merge tool with miRBase (release 21), allowing for a maximum of 1 mismatch. Differential expression analysis was performed using the Empirical Analysis of DGE tool with false discovery rate (FDR) correction.

#### 2.5.4 Total RNA composition

Total composition of known small RNAs was calculated in CLC Genomics by performing annotate and merge on known small RNAs extracted from amel_OGSv3.2. To determine the total reads mapping to *A*. *mellifera*, DWV and BQCV genomes, trimmed reads were mapped first to the *A. mellifera* genome (amel_OGSv3.2) using the map reads to reference tool, unmapped reads were subsequently mapped to DWV and BQCV.

A final check of the unmapped reads was performed by generating a *de novo* assembly for each sample using Megahit (Li et al., 2015). As well as generating contigs representing DWV and BQCV genomes, multiple contigs were assembled that showed near 100% identity to *A. mellifera* ribosomal RNA in BLASTn searches. Unmapped reads from each sample were mapped back to the assembled rRNA contigs using Bowtie2 (Langmead and Salzberg, 2012), to assign any remaining ribosomal reads from the unmapped reads to their correct category.

## 3. RESULTS AND DISCUSSION

Our main objective was to determine if honey bees naïve to both *V. destructor* and DWV differ in their immune response when exposed to DWV-A and DWV-B. Injecting honey bees with buffer is known to activate covert viral infections (Anderson and Gibbs, 1988; Dubois et al., 2019) and is likely to trigger an immune response to wounding (Kuster et al., 2014). We therefore included two control groups, an unmanipulated pupae control and a buffer-injected pupae control. By comparing the immune response of DWV or buffer injected pupae (BC) to unmanipulated pupae (UC), we were able to separate the effect of DWV genotypes from the effect of experimental injection.

### 3.1 Up regulation of immune genes in DWV infected pupae

Of the 19 immune genes examined, the majority [13 genes, excluding *domeless* and the antimicrobial peptides (AMPs)] were expressed at higher levels in pupae from 48-72 HPI onwards (Figure 1; Table S2). Some genes (*dicer*, *argonaute-2*, *relish*, *malvolio*, *hopscotch*) were strongly up-regulated from 48 HPI in DWV injected pupae. Gene up-regulation coincided with the accumulation of DWV. Loads of both DWV genotypes exceeded 1 × 10^7^ (relative to *Actin*) at 48 HPI, and exceeded 1 × 10^8^ and 1 × 10^9^ in DWV-A and DWV-B pupae by 96 HPI, respectively [Figure 1A; (Norton et al., 2020)].

**Figure 1.**
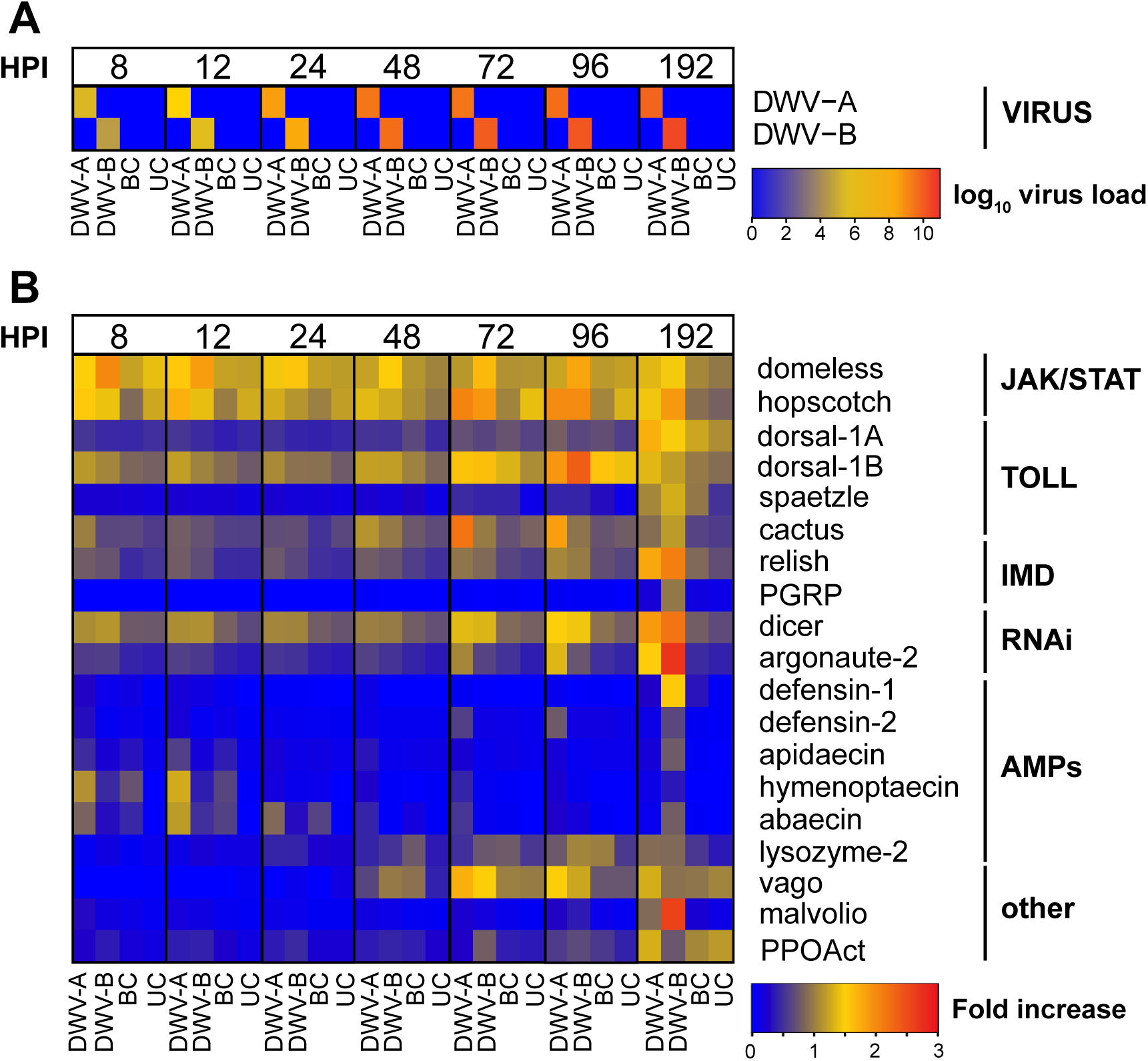
A. Heat map A shows the median log_10_ viral load of DWV in pupae injected with either DWV-A, DWV-B, or buffer (BC), and unmanipulated pupae (UC) at 8 to 192 hours-post injection. Each row illustrates the level of DWV-A and DWV-B in pupae, and each column corresponds to a different treatment group at each timepoint. The scale bars indicate median log_10_ virus load relative to *Actin.* **B.** Heat map B displays the relative expression of 19 immune genes in the same pupae, where each row corresponds to a gene. The scale bars indicate the median increase in gene expression relative to two housekeeping genes (*Actin* and *RpS5*). The immune pathways corresponding to each gene are indicated on the far right.

RNAi is the major antiviral mechanism in insects (Gammon and Mello, 2015). It was therefore expected that honey bees would mount an RNAi response when injected with DWV. The expression of both *dicer* and *argonaute-2* increased in parallel with DWV viral loads in pupae over time. *Dicer* was the only gene that was up-regulated in DWV injected pupae compared to both control groups at all time points (Figure 2). Median expression of *dicer* was approximately 0.5-fold higher in DWV injected pupae compared to the UC group from 8-48 HPI, and increased 0.6-2.8 fold between 72-192 HPI (Table S3). The expression of *argonaute-2* ranged from 0.4 to 7.1-fold higher in DWV injected pupae compared to the UC group between 8-192 HPI, but did not differ to the BC group at 48 HPI and 24-48 HPI in DWV-A and DWV-B pupae, respectively (Figure 2; Table S4). In a previous study, neither *dicer* or *argonaute-2* were found to be differentially expressed in purple-eyed pupae parasitised by *V*. *destructor* (approximately equivalent to our 24 HPI), despite detecting a siRNA response (Ryabov et al., 2014). However, our results show that the RNAi response is time-dependent with increased expression associated with accumulating viral loads over time.

**Figure 2.**
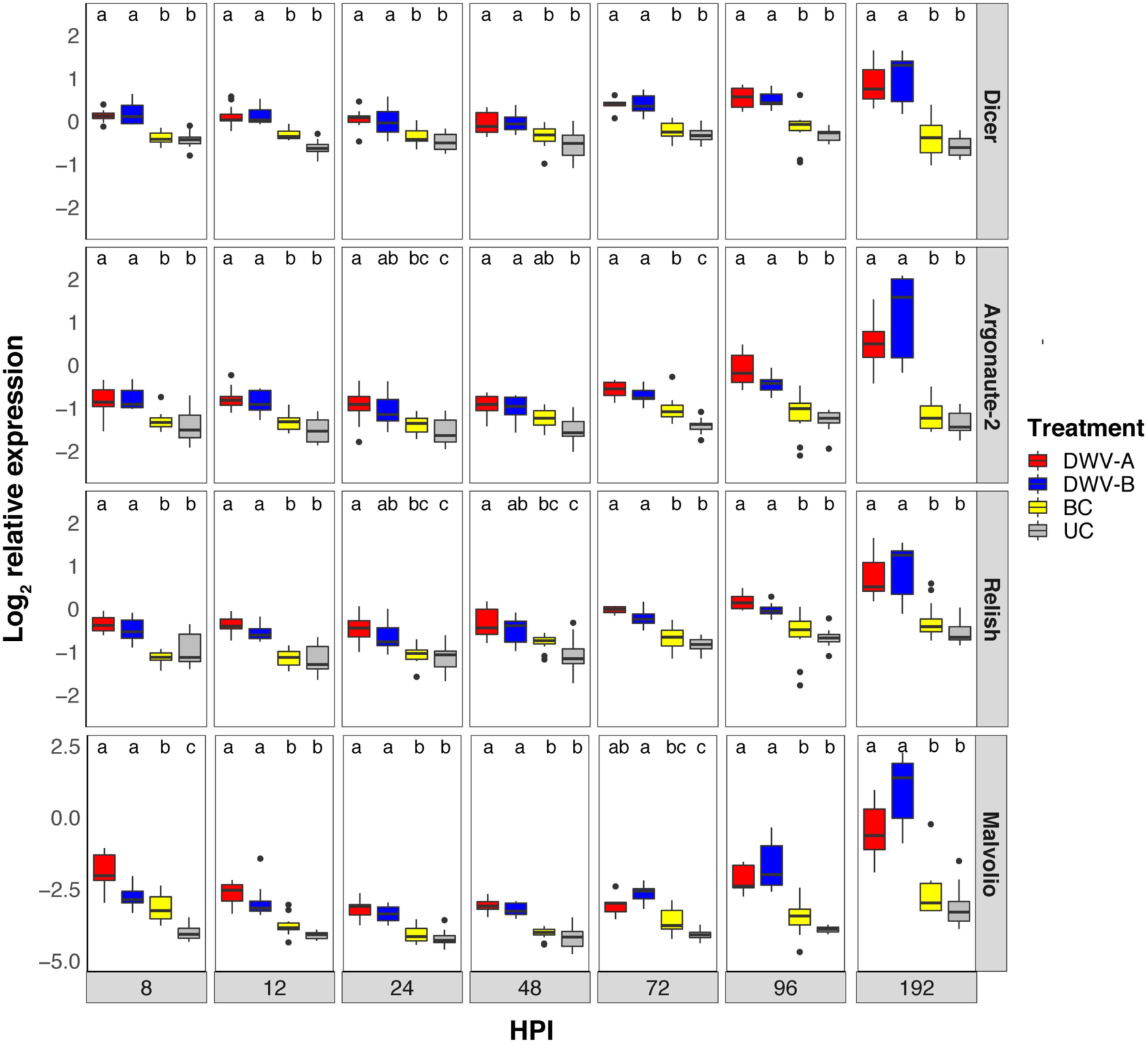
Log_2_ relative expression of *dicer*, *argonaute-2*, *relish* and *malvolio* in honey bee pupae. Treatment groups that do not share a common letter differ at *p* < 0.05 at each hour post-injection (HPI).

*Relish* is a transcription factor in the Imd pathway, which upon cleavage, leads to the production of antimicrobial peptides (AMPs) *abaecin* and *hymenoptaecin* (Schlüns and Crozier, 2007). Similarly to *dicer* and *argonaute-2,* we found increased expression of *relish* in DWV injected pupae, which differed from both controls at all time points, except DWV-B and BC pupae between 24-48 HPI (Figure 2; Table S5). The highest expression of *relish* occurred at 192 HPI, which was 1.3-2.8-fold higher than in UC pupae. A previous study found an association between high DWV loads and mite infestation with increased expression of *relish* in honey bee colonies, suggesting that *relish* is an immunological marker of DWV-*Varroa* parasitism (Barroso-Arévalo et al. (2019b). Our results indicate that DWV alone (in the absence of *V*. *destructor*) triggers up-regulation of *relish*.

The metal ion (Mn^2+^, Fe^2+^ and Cu^2+^) transporter *malvolio* was up-regulated in DWV injected pupae compared to both control groups at all time points, excluding DWV-B compared to BC pupae at 8 HPI (*Z* = 1.21, *p* = 0.226) and DWV-A relative to BC pupae at 72 HPI (*Z* = 1.67, *p* = 0.114) (Figure 2; Table S6). Expression was 1.1-5.3 and 0.8-24.7-fold higher in DWV-A and DWV-B pupae, respectively, compared to the UC group. We found the highest expression at 192 HPI, yet due to the variability between samples, differences between DWV-A and DWV-B pupae were not statistically significant (*Z* = –1.36, *p* = 0.207). *Malvolio* is involved in the sucrose responsiveness of insects, and thus affects foraging behaviour (Søvik et al., 2017). Foraging honey bees, who consume a carbohydrate rich diet, express higher levels of *malvolio* compared to nurse bees who consume higher amounts of protein. Feeding manganese to newly emerged honey bees results in increased expression of *malvolio* and premature foraging (Ben-Shahar et al., 2004), suggesting a cause and effect relationship. *Malvolio* is also up-regulated in mite-parasitised honey bees infected with DWV (Alaux et al., 2011; Zanni et al., 2017). Interestingly, experimental injection of DWV also induces precocious foraging in honey bees (Benaets et al., 2017; Natsopoulou et al., 2016). Taken together, increased expression of *malvolio* could potentially explain premature foraging in DWV infected honey bees.

The transmembrane receptor *domeless* and the kinase *hopscotch* are both part of the JAK/STAT pathway, associated with responses to immune challenge and septic injury in mosquitoes and *Drosophila* (Lematire and Hoffman 2007). We found increased expression of *domeless* in DWV-B injected pupae where expression was approximately 0.5-fold higher compared to both controls, excluding 24 HPI (Figure S1; Table S7). However, the highest expression was observed at 8-12 HPI when DWV-B viral loads were < 1 × 10^6^ (Norton et al., 2020). This suggests that increased expression in DWV-B pupae was independent of viral replication. In contrast, *domeless* expression in DWV-A pupae was equal to both control groups, excluding 12 and 192 HPI. In *Drosophila*, *hopscotch* is activated by *domeless* and subsequent phosphorylation creates STAT92E binding sites, which induces the transcription of genes involved in antiviral activity (Merkling and van Rij, 2013; Öhlund et al., 2019). Despite limited differences in *domeless* expression in our DWV injected bees compared to the controls, we detected 0.4 to 1.4-fold higher expression of *hopscotch* in DWV injected bees compared to both control groups at 96-192 HPI (Figure S1; Table S8). In *Drosophila*, up-regulation of *hopscotch* induces the transcription of *vir-1*, which counters the replication of *Drosophila C virus* (Dostert et al., 2005). It is unclear whether the JAK/STAT pathway plays a role in antiviral defence against DWV in honey bees. Previous studies have found that *domeless* and *hopscotch* are up-regulated in DWV infected bees parasitised by *V*. *destructor* (Tesovnik et al., 2019; Zhao et al., 2019), while others found the opposite (Khongphinitbunjong et al., 2015; Ryabov et al., 2014).

### 3.2 Toll pathway response to DWV

It has previously been suggested that the Toll pathway, particularly expression of NF-κB factor *dorsal-1A,* is a key element in regulating the honey bee immune response against DWV (Nazzi et al., 2012; Nazzi and Pennacchio, 2018). *Dorsal-1* has two splice variants (*1A and 1B*), which are primarily expressed in the fat body tissue of honey bees, and *dorsal-1A* regulates the expression of *defensin-1* (AMP) (Lourenco et al., 2018). *Cactus* is a NF-κB inhibitor, which must be degraded to allow nuclear translocation of *dorsal* (Zuo et al., 2016). Increased *cactus* expression in DWV-infected adult bees parasitised by *V*. *destructor* has been associated with the down-regulation of *dorsal* (Zanni et al., 2017). Thus, we expected to find increased expression of *cactus* coupled with the down-regulation of *dorsal-1A* in our DWV injected pupae. However, our data showed that the expression of *dorsal-1A* (Figure 3; Table S9) and *dorsal-1B* (Figure S2; Table S10) in DWV injected pupae was very similar to both control groups with only minor differences between treatment groups over time. The expression of *cactus* was also generally very similar between DWV injected pupae and both controls (Figure 3). We did, however, find elevated expression of *cactus* in DWV injected pupae at 48 and 192 HPI (Table S11), where median expression was 0.6-1.2 fold higher than in the UC group. However, this increase was not associated with any notable changes in *dorsal-1A* expression. As RNAi knockdown of *dorsal-1A* resulted in increased DWV loads, Nazzi et al. (2012) suggested that the transcription factor plays in important role in antiviral defence. *Dorsal-1A* was also down-regulated in DWV infected bees but mite parasitism did not have an effect on expression (Nazzi et al., 2012), even though *V*. *destructor* feeds on the fat bodies of honey bees (Ramsey et al., 2019). Annoscia et al. (2019) later found that experimental removal of honey bee haemolymph reduces *dorsal-1A* expression, although this was also associated with increased accumulation of DWV. Our results suggest that either *dorsal-1A* expression is only down-regulated in honey bees that have been pre-exposed to DWV, or that DWV alone (in the absence of *V*. *destructor* feeding) does not down-regulate *dorsal-1A*. Further, *spaetzle* was not down-regulated in our DWV injected bees, in contrast to previous findings in DWV infected and mite infested pupae (Khongphinitbunjong et al., 2015; Ryabov et al., 2014). Instead, we found that the expression of *spaetzle* was equally elevated in DWV and BC injected pupae (Figure 3; Table S12), suggesting that the up-regulation of *spaetzle* was likely attributed to wounding during injection rather than DWV infection.

**Figure 3.**
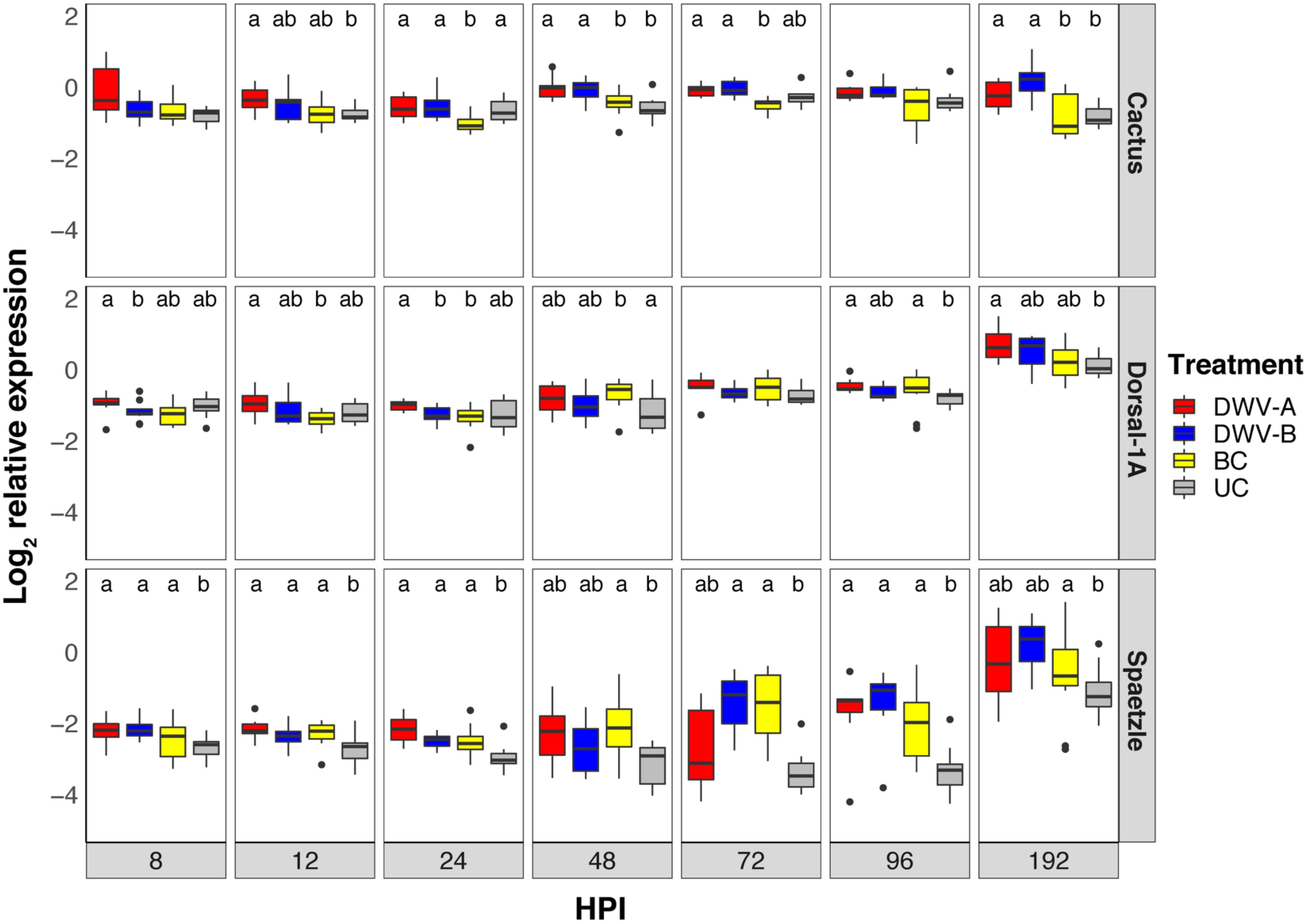
Log_2_ relative expression of *cactus*, *dorsal-1A* and *spaetzle* in honey bee pupae. Treatment groups that do not share a common letter differ at *p* < 0.05 at each hour post-injection (HPI). No letters within a given time point indicates that no significant differences were detected between treatment groups.

### 3.3 Up-regulation of AMPs at early time points due to wounding

The majority of previous studies have found an increase in expression of antimicrobial peptides (AMPs) in DWV infected bees (Table 1), both when experimentally injected (Ryabov et al., 2016) and mite parasitised (Khongphinitbunjong et al., 2015; Kuster et al., 2014; Zanni et al., 2017; Zhao et al., 2019). We also observed increased AMP expression, but only at specific time points. Within the first ∼ 48 hours of infection, expression of AMPs was often equal in the BC and DWV injected pupae (Tables S13-S17). This was most evident in the expression of *abaecin, hymenoptaecin*, and *defensin-1* (Figure 4), where up-regulation from 8 to ∼48 HPI was likely due to wounding. However, we did find an approximate 1-fold increase in expression of *abaecin* in DWV-A injected pupae at 8 (*Z* = 2.23, *p* = 0.031) and 12 HPI (*Z* = 2.07, *p* = 0.046) (Figure 4) and *defensin-2* at 8 HPI (*Z* = 2.71, *p* = 0.013) relative to the BC group (Figure S3). Expression of *abaecin, hymenoptaecin*, and *defensin-1* in the BC pupae was more similar to the UC group in the later time points. *Hymenoptaecin* was up-regulated in both DWV-A and DWV-B injected bees from 72-192 HPI, which might be explained by the increased expression of *relish* at the same time points. We found a similar increase for *abaecin* at 96 HPI, but only DWV-B injected pupae differed from the BC group at 192 HPI. The down-regulation of *defensin-1*, which is under the control of *dorsal-1A* (Lourenco et al., 2018), was associated with increased DWV loads and mite infestation in a previous study (Barroso-Arévalo et al., 2019b). In contrast, we found no evidence to indicate that *defensin-1* is down-regulated as a result of DWV infection. Conversely, we often found that AMP expression was higher in DWV-B injected pupae at 192 HPI compared to the BC group, including *defensin-1*, whereas expression in DWV-A pupae often did not differ from the BC group at 192 HPI (Figure 4; Figure S3).

**Figure 4.**
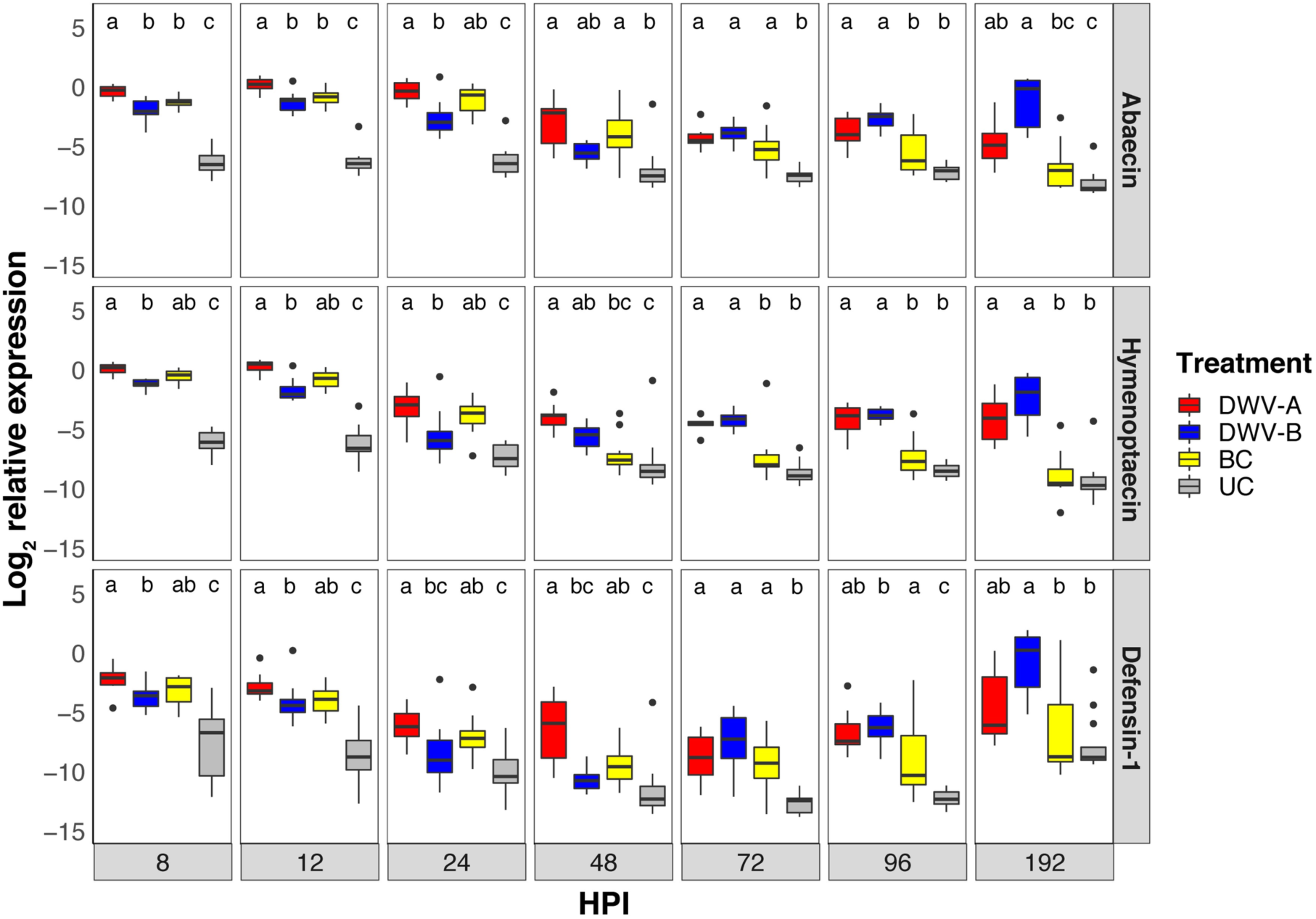
Log2 relative expression of *abaecin, hymenoptaecin* and *defensin-1* in honey bee pupae. Treatment groups that do not share a common letter differ at *p* < 0.05 at each hour post-injection (HPI).

**Figure 5.**
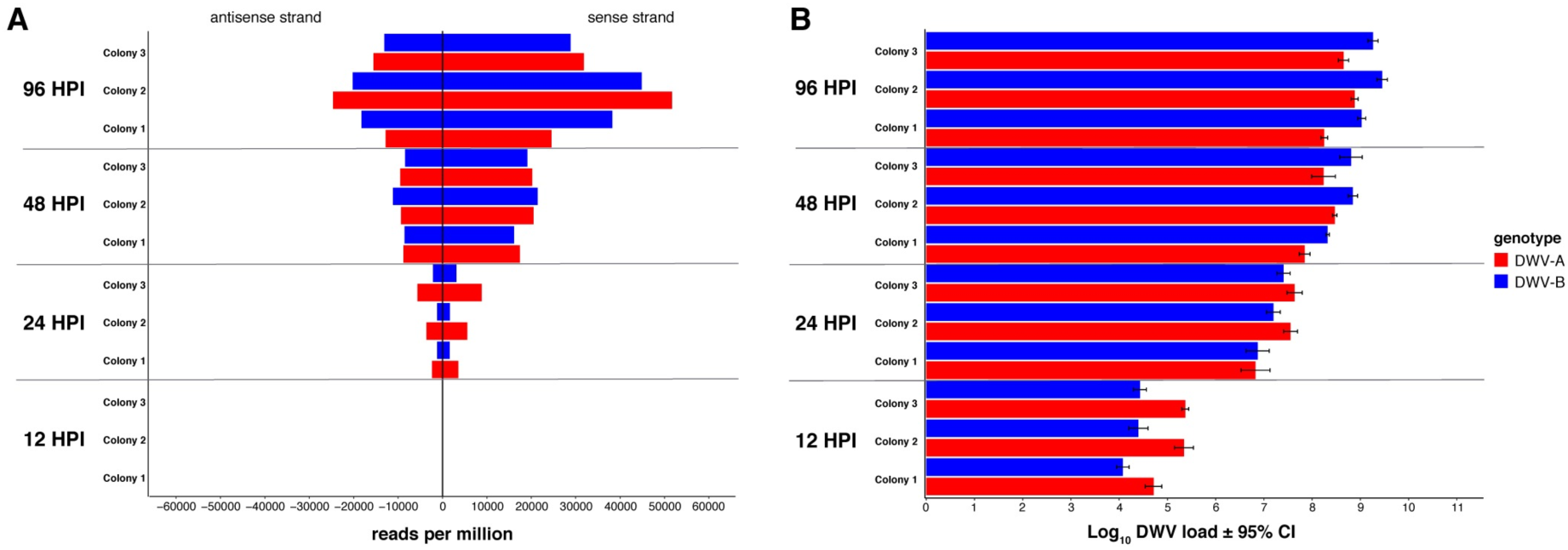
Small RNA analysis in honey bee pupae. (A) The total sense and antisense vsiRNA (21-23 nt) reads mapping to DWV-A (red) and DWV-B (blue), in four pooled honey bee pupae per time point and colony. Reads at 12 HPI are not visible at this scale. (B) The mean DWV loads ± 95% CI of four pupae per treatment and colony at 12, 24, 48 and 96 hours post-injection. Viral loads are relative to housekeeping gene *Actin* and were previously published in Norton et al. (2020), but are presented here to illustrate the relationship between viral accumulation and siRNA response to DWV-A and DWV-B.

### 3.4 Differential expression in response to DWV-A and DWV-B

We find very little evidence to indicate that honey bees respond differently to DWV-A and DWV-B. If differences in immune response were biologically significant, we would have expected a consistent difference in gene expression over time, or within a given immune pathway. However, this was not the case. We only found differences at isolated time points, where the expression in pupae exposed to one genotype was also different to both controls. When we did see a difference between two genotypes, this was often associated with higher expression in DWV-A injected pupae at early time points [*malvolio:* (8 HPI, 0.8-fold; *Z* = 2.10, *p* = 0.043); *defensin-2:* (8 HPI, 1-fold; *Z* = 2.76, *p* = 0.017); and *abaecin:* (8 HPI, 2.4-fold *Z* = 3.24, *p* = 0.002; and 12 HPI, 1.6-fold; *Z* = 2.98, *p* = 0.006)]. DWV-A accumulates to higher loads than DWV-B within the first 12 hours post-injection (Figure 1A and 5B) so it is possible that these differences are associated with faster replication of DWV-A during the initial stage of infection. In contrast, we found that gene expression was often elevated in DWV-B injected pupae at later time-points (192 HPI), although this was only statistically different from DWV-A for *PGRP-S2* (Toll) (5-fold; *Z* = –2.36, *p* = 0.036) (Figure S2; Table S18), and *defensin-2* (9-fold; *Z* = –2.40, *p* = 0.033) (Figure S3), other than increased expression of *domeless* at 8 HPI (0.3-fold; *Z* = –2.67, *p* = 0.015) and *PPOact* at 72 HPI (0.9-fold; *Z* = –2.71, *p* = 0.013) (Figure S4; Table S19-S21). This is consistent with elevated viral loads of DWV-B at later time points [(Figure 1A and 5B; Norton et al. (2020)].

### 3.5 Small RNA analysis

#### 3.5.1 siRNA response to DWV genotypes

Our expression analysis showed that RNAi pathway genes *dicer* and *argonaute-2* are clearly up-regulated in DWV injected pupae (Figure 2). Previously, we found that DWV-B accumulated to 5 to 10-fold higher loads than DWV-A (Figure 1A; Norton et al., 2020) from 48-192 HPI, suggesting that DWV-B may be better able to evade RNAi degradation compared to DWV-A. Hence, we investigated the small RNA profiles of DWV injected pupae to determine if honey bees elicit a different siRNA response to either genotype. We analysed four pooled DWV-A and DWV-B injected pupae per colony at 12, 24, 48 and 96 HPI. The number of reads mapping to DWV increased over time, ranging from < 1.6% of the total reads for each sample at 12 and 24 HPI, to 3-4% at 48 HPI and 4.4-8.5% at 96 HPI. We examined the read size profiles of each sample and saw that a clear antiviral small RNA response was elicited in pupae injected with DWV-A and DWV-B. A signature *dicer* response was observed in pupae collected at 24, 48, and 96 HPI, whereby vsiRNA reads (mapping to DWV-A and DWV-B genomes) were predominantly 22 nt in length, followed by 21 and 23 nt. Reads showed a 1.3-2.3-fold bias towards the sense orientation (Figure S5.A), consistent with previous observations of DWV infected honey bees (Chejanovsky et al., 2014; Ryabov et al., 2014; Wang et al., 2013). At 12 HPI there were higher counts of 15-16 nt fragments followed by less abundant 21-23 nt fragments. We found that the vsiRNAs (21-23 nt) in both DWV-A and DWV-B injected pupae increased over time, with counts ranging from 8.4-182/million reads at 12 HPI to 37400-76300/million reads at 96 HPI (Figure 5A). In agreement with Ryabov et al. (2014), we found that vsiRNA response in pupae strongly correlated with viral load of DWV-A (*r_s_* = 0.98, *p* < 0.0001) and DWV-B (*r_s_* = 0.99, *p* < 0.0001) (Figure 5B). However, we found no difference in the vsiRNA response between pupae injected with DWV-A or DWV-B (χ^2^= 0.27, df = 1, *p* = 0.6033) (Figure S5.B), or between the three colonies (χ^2^= 0.455, df = 2, *p* = 0.7965). This suggests that previously observed differences in viral accumulation (Norton et al., 2020), are likely explained by replicative characteristics of the DWV genotypes themselves rather than RNAi-mediated degradation by the host. We also found that the vsiRNAs were randomly distributed across both DWV genomes, yet we consistently detected a large peak starting at nucleotide position 8180 in DWV-A injected pupae (Figure S5.C) that was absent in DWV-B injected bees (Figure S5.D). Interestingly, a peak at a similar position was observed in Israeli colonies (Chejanovsky et al., 2014). The strong peak could be due to active targeting by RNAi or could be indicative of a viral miRNA (Bruscella et al., 2017; Hussain et al., 2011), however further analysis is required to establish this.

#### 3.5.2 piRNA response

The PIWI-interacting RNA (piRNA) response (RNAi) is known to play a role in germline protection against transposable elements in *Drosophila* (Czech and Hannon, 2016). piRNAs typically range from 26-31 nt in length and are produced independently of *dicer*, from single-stranded RNA (Hirakata and Siomi, 2016). The piRNA response also plays a role in antiviral defence against *Chikungunya* and *Dengue virus* in *Aedes aegypti* and *Ae. albopictus* mosquitos (Goic et al., 2016; Miesen et al., 2016). We analysed the larger sized sRNA reads (26-31 nt) that would be indicative of viral piRNAs but did not detect any obvious peaks. We did, however, detect a low abundance of 26-31 nt fragments in all samples, which increased over time. Counts ranged from ∼ 0-122/million reads at 12-24 HPI and ∼22-200000/million reads at 48-96 HPI, with a strong bias towards the sense orientation. To confirm that viral piRNAs were not produced at low levels, the 26-31 nt reads were analysed for the ‘ping-pong’ (1U/10A) signature associated with piRNA amplification (Czech and Hannon, 2016). Our results showed slightly (< 40%) elevated frequencies of uridine at the 5’ end, however we found no evidence of adenine bias at position 10 (Figure S6). We also observed elevated frequencies of uridine at the 3’ end. Overall, the data suggested that the 26-31 nt reads were not piRNAs, but products of random degradation.

#### 3.5.3 miRNA profiles

We also mapped the small RNA reads to the honey bee (*Apis mellifera*) genome and determined the composition of small RNA subtypes within each sample (Figure 6). There was some variability between the samples. The samples were predominantly made up of rRNA (15-58%), followed by miRNA (13-38%), tRNA (2-13%) and mRNA (0.8-2%). We then analysed the miRNA profiles. We found a small number of differentially expressed miRNAs (FDR-corrected *P* < 0.01) between pupae at 24, 48 and 96 compared to the 12 HPI pupae. The number of differentially expressed miRNAs increased over time in both DWV-A (0 to 28) and DWV-B (3 to 38) injected pupae. However, as all of our libraries were generated from DWV-A or DWV-B injected samples, we were unable to distinguish whether this was associated with viral infection or physiological changes at different time points during pupation. We subsequently compared the miRNAs expressed between DWV-A and DWV-B injected pupae at each specific time point, and found no differentially expressed miRNAs at 12-48 HPI, but found three at 96 HPI. Mir-3720 (FDR corrected *p* < 0.0001) and mir-6052 (*p* < 0.0001) were expressed 6.7 and 11.5-fold higher in the DWV-A injected pupae, respectively, whereas mir-279d was expressed 1.6-fold higher in DWV-B injected pupae (*p* = 0.004). However, we cannot exclude the possibility that differential miRNA expression was associated with *Black queen cell virus* [BQCV (see below)]. All three miRNAs have previously been identified in honey bees (Chen et al., 2010; Liu et al., 2012; Macedo et al., 2016; Qin et al., 2014), however their function is currently unknown.

**Figure 6.**
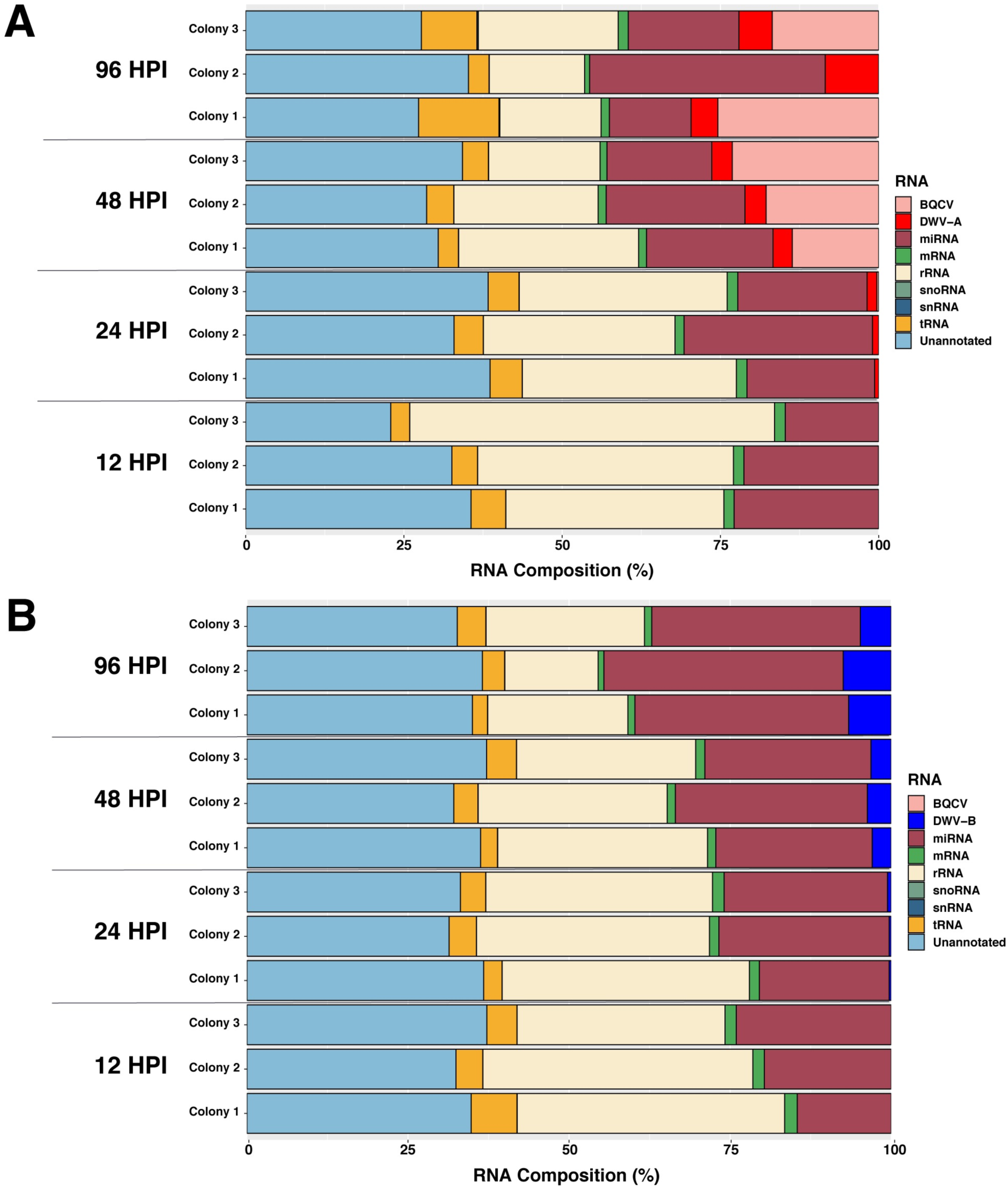
Percentage composition of RNA in (A) DWV-A and (B) DWV-B injected pupae. DWV-A and DWV-B at 12 HPI, and snoRNA and snRNA at all time points constituted < 1% of total RNAs.

#### 3.5.4 siRNA response to BQCV

We additionally performed a *de novo* assembly of the unmapped reads and used the resulting contigs to perform a BLAST search. We detected BQCV in five of our DWV-A injected samples (Figure 6A), as expected based on our previous detection of BQCV in DWV-A injected pupae (Norton et al., 2020). We detected very low reads for BQCV at 12 (< 0.001%) and 24 HPI (< 0.3%), consistent with our previous qPCR results where BQCV was undetected or ≤ 1 × 10^2^ (relative to *Actin*) at these time-points. The number of small RNA reads aligning to BQCV ranged from 14-23% at 48 HPI and 0-25% at 96 HPI, as BQCV was not detected in colony 2 at 96 HPI. BQCV was not detected in DWV-B injected pupae (Figure 6B), in agreement with our previous qPCR results (Norton et al., 2020).

We analysed the small RNA profiles of the reads aligning to BQCV and found that samples containing both viruses exhibited a stronger siRNA response to BQCV compared to DWV. Again, the reads were predominantly 22 nt in length, with a 1.5 to 2-fold bias to the sense orientation (Figure S7.A). However, this time the vsiRNAs (21-23 nt) ranged from 1200000-3300000/million reads (Figure S8.A), which exceeded the siRNA response to DWV by approximately two orders of magnitude. This is in contrast to the viral loads observed in the same samples, where high BQCV loads were ∼5 to 10-fold lower than DWV-A loads in individuals containing both viruses [Figure 1A and S8.B; Norton et al. (2020)]. BQCV loads were highly variable between pupae (Figure S8.B), due to the majority of pupae per pooled group harbouring low BQCV loads < 3 × 10^4^, and there was no correlation between BQCV load and small RNA response (*r_s_* = 0.2; *P* = 0.7471). We found that vsiRNAs were randomly distributed across the BQCV genome (Figure S7.B). BQCV has been found to exist at covert levels in our population of honey bees (Remnant et al., 2019) and is common pathogen of honey bees worldwide (McMahon et al., 2015; Murray et al., 2018; Roberts et al., 2017). Yet, observing such a large siRNA response to BQCV was surprising, considering so few pupae harboured high BQCV loads. Our results suggest that BQCV may elicit a stronger response than DWV because it is a more damaging virus at the pupal stage compared to DWV, which does not lead to high pupal mortality (Remnant et al., 2019), or that our population of bees may be better able to target BQCV due to pre-exposure to the virus. Alternatively, our results may indicate that DWV is comparatively better at evading siRNA degradation. Some RNA viruses are known to evade RNAi by encoding suppressors (VSR). These VSRs can inhibit RNAi through a number of different mechanisms, such as binding long dsRNA or double stranded siRNAs thereby inhibiting *dicer* cleavage or RISC loading (Nayak et al., 2010; Sullivan and Ganem, 2005; Van Rij et al., 2006), or may interfere with RNAi by directly interacting with *dicer*, *argonaute* or other siRNA processes (Nayak et al., 2010; Qi et al., 2012; Singh et al., 2009). It is unknown whether DWV uses VSRs to evade RNAi, however a putative VSR has been identified in *Israeli acute paralysis virus* (Chen et al., 2014), a dicistrovirus honey bee pathogen related to BQCV.

## 4. CONCLUSION

Previous studies have found that the two main DWV genotypes, A and B, affect honey bees differently (Dubois et al., 2019; Gisder et al., 2018; McMahon et al., 2016; Norton et al., 2020; Tehel et al., 2019), although results differ between life stages and honey bee populations, leaving our understanding of DWV genotype virulence unclear. Over time, the prevalence of the two main DWV genotypes has changed, where DWV-B has increased at the expense of DWV-A (Kevill et al., 2019; Ryabov et al., 2017). The elevated prevalence of DWV-B suggests that the two genotypes may interact differentially with the honey bee immune system (Mordecai et al., 2016). Here we investigated the effect the two genotypes have in isolation, and in the same *V*. *destructor* and DWV-naïve population. Overall, we found no evidence to indicate that the honey bee’s immune system responds to either genotype differently. We only detected isolated differences in gene expression, and these may be attributed to differences in viral loads. This suggests that global changes in DWV-B prevalence are not explained by an immunologically different response in honey bee hosts. Our work highlights that the siRNA pathway is up-regulated in response to DWV infection, but that the host degradation response does not inhibit the virus from accumulating to high loads. We found that differences in viral accumulation between the two DWV genotypes are not associated with differential RNAi expression. Further, our results showed that the siRNA response towards BQCV far exceeded the response to DWV, implying that the high pathogenicity of BQCV elicits a stronger immune response or that DWV is comparatively better at evading antiviral host defences. Future studies are required to investigate what mechanisms DWV may utilise to evade RNAi defence by honey bee hosts.

*Deformed wing virus*, in association with the parasitic mite *V*. *destructor*, has been associated with the death of millions of honey bee colonies worldwide (Dainat et al., 2012; Martin, 2001; Schroeder and Martin, 2012). *V*. *destructor* was thought to have an immunosuppressive effect on honey bees (Yang and Cox-Foster, 2005). Recent studies conclude that mite feeding upon fat bodies directly weakens bees and disrupts their immune response (Annoscia et al., 2019; Kuster et al., 2014; Ramsey et al., 2019), particularly as fat bodies are a major site of immune and metabolic function (Arrese and Soulages, 2010). Indeed, Annoscia et al. (2019) showed that experimental removal of haemolymph (likely containing fat body tissue) resulted in the down-regulation of *dorsal-1A* and parallel increase of DWV loads naturally present within honey bees. The authors concluded that mite feeding disrupts the balance between honey bee immune response leading to increased viral replication. The fact that DWV is now found in virtually all honey bee populations worldwide (Martin and Brettell, 2019), makes it challenging to distinguish whether the down regulation of *dorsal-1A* was attributable to viral replication as previously suggested (Nazzi et al., 2012) or a result of haemolymph removal. Our results show that actively replicating DWV, in the absence of *V*. *destructor*, does not down-regulate *dorsal-1A*, supporting previous conclusions that mite feeding on bee tissue disrupts immune function (Annoscia et al., 2019; Kuster et al., 2014; Ramsey et al., 2019).

## DATA AVAILABILITY

*De novo* assembly of the BQCV genome sequence has been deposited to GenBank (accession number MW390818). Supporting data files including raw small RNA reads are available at https://doi.org/10.6084/m9.figshare.13473696

## ACKNOWLEDGEMENTS

We thank Boris Yagound and Nicholas M. A. Smith for their advice with the bioinformatics analyses, and acknowledge the Sydney Informatics Hub and The University of Sydney’s high performance computing cluster Artemis for providing the high performance computing resources that have contributed to the research results reported within this paper.

## AUTHOR CONTRIBUTIONS

AMN, EJR and MB conceived the study. AMN conducted the experimental work. AMN and GB carried out molecular laboratory work. GB conducted the small RNA library preparation. AMN analysed the qPCR and statistical data. AMN and EJR analysed the siRNA data. AA conducted the miRNA analysis. OW analysed the piRNA data. AA and EJR determined the total composition of known small RNAs. AMN, MB and EJR contributed to the writing of the manuscript. All authors gave final approval for publication.

