## Supplementary Figures S1-S8 for "Deformed wing virus genotypes A and B do not elicit immunologically different responses in naïve honey bee hosts"

This file contains:

- Supporting Figures: S1-S8

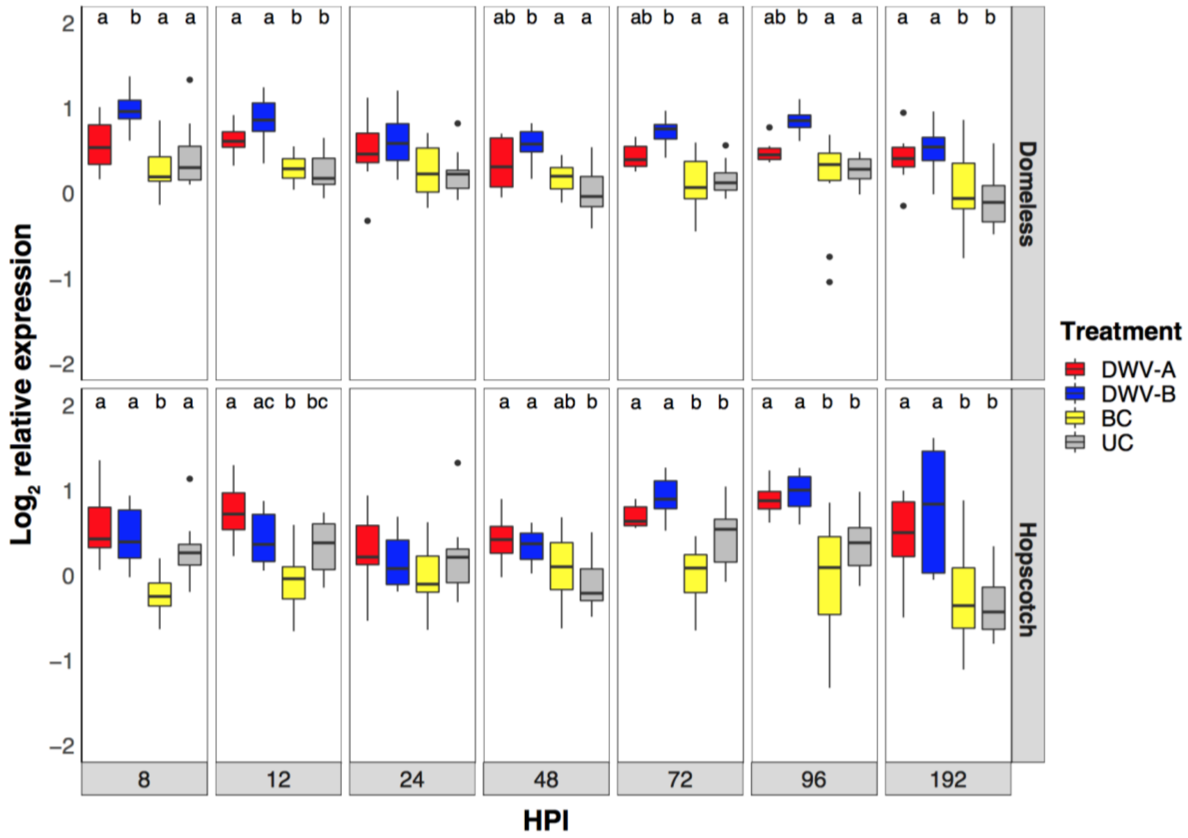


**Figure S1.** $\text{Log}_2$ relative expression of JAK/STAT pathway genes *domeless* and *hopscotch* in honey bee pupae. Treatment groups that do not share a common letter differ at $p < 0.05$ at each hour post-injection (HPI). No letters above a given time point indicates that no significant differences were detected between treatment groups.

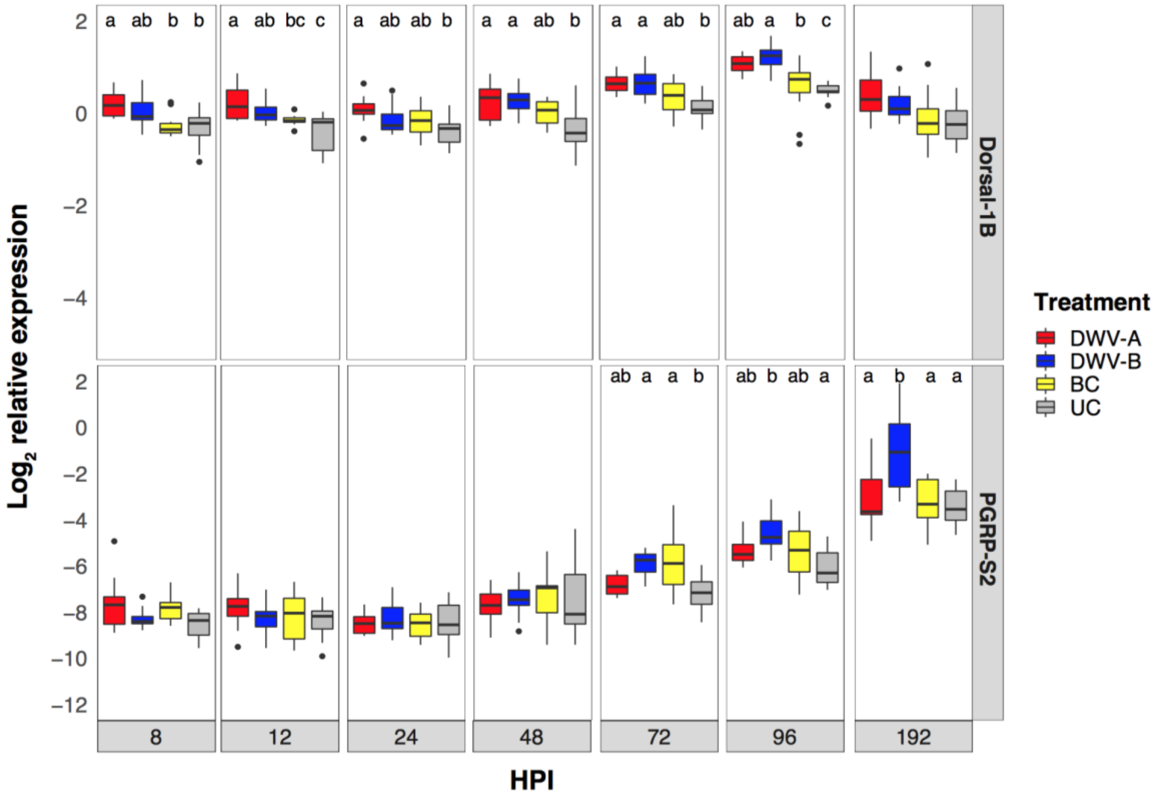


**Figure S2**. $Log_2$ relative expression of Toll pathway genes *dorsal-1B* and *PGRP-2* in honey bee pupae. Treatment groups that do not share a common letter differ at $p < 0.05$ at each hour post-injection (HPI). No letters above a given time point indicates that no significant differences were detected between treatment groups.

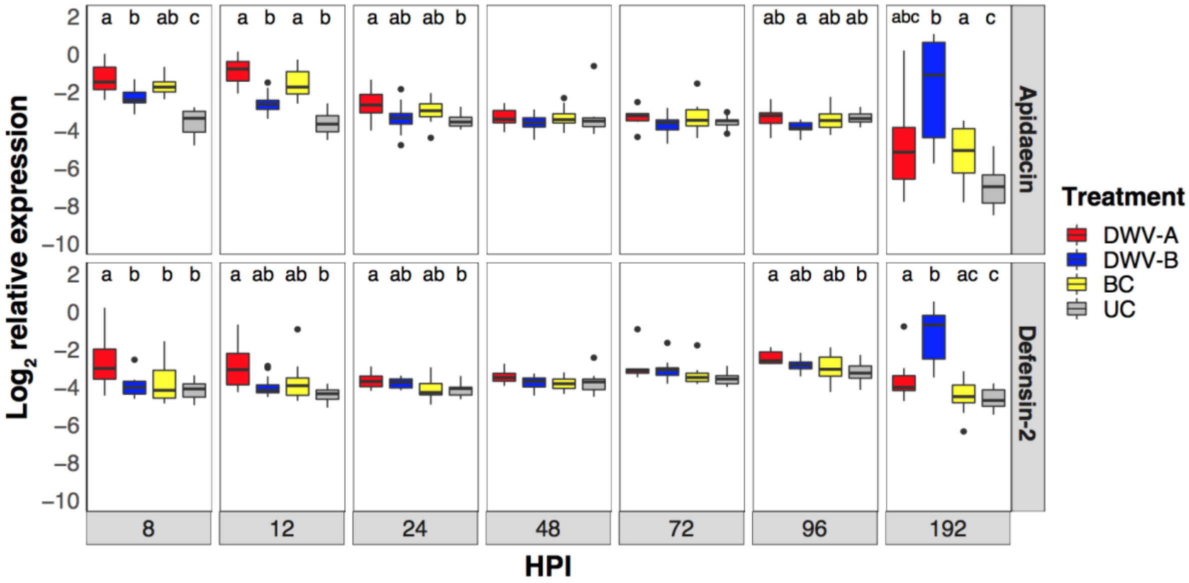


**Figure S3.** $Log_2$ relative expression of AMP genes *apidaecin* and *defensin-2* in honey bee pupae. Treatment groups that do not share a common letter differ at $p < 0.05$ at each hour post-injection (HPI). No letters above a given time point indicates that no significant differences were detected between treatment groups.

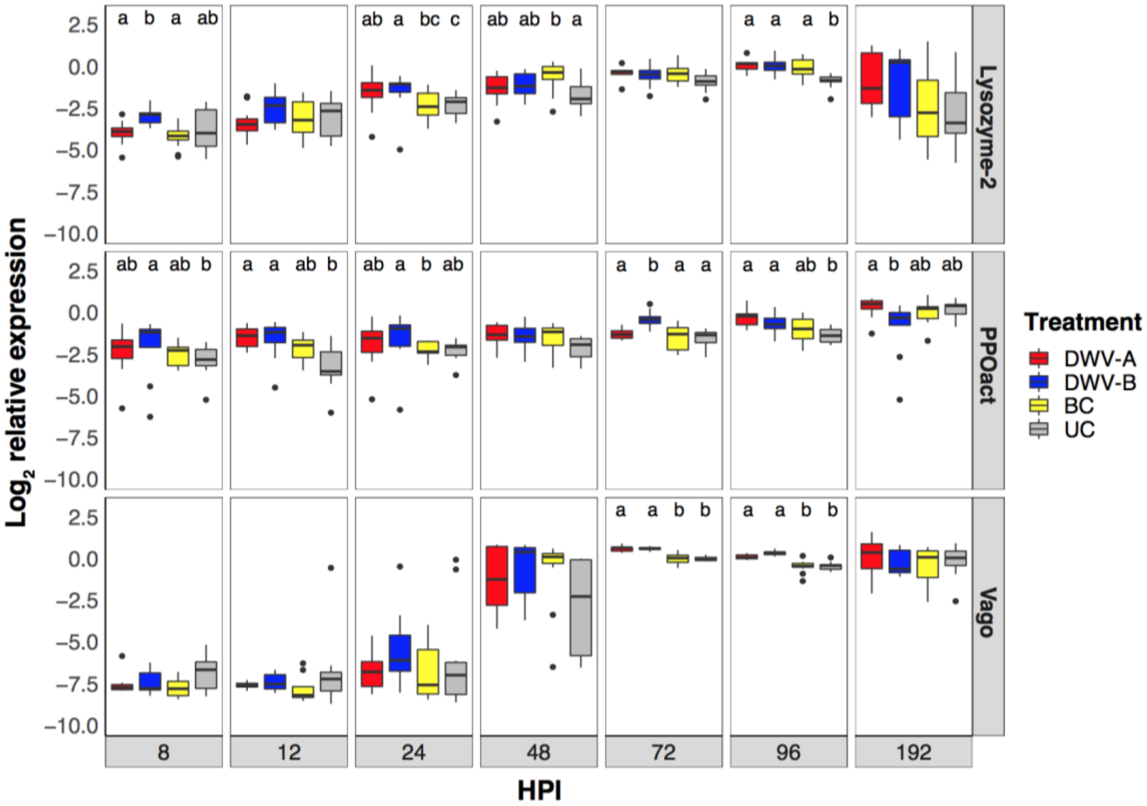


**Figure S4.** Log2 relative expression of genes *lysozyme-2*, *PPOact* and *vago* in honey bee pupae. Treatment groups that do not share a common letter differ at $p < 0.05$ at each hour post-injection (HPI). No letters above a given time point indicates that no significant differences were detected between treatment groups.

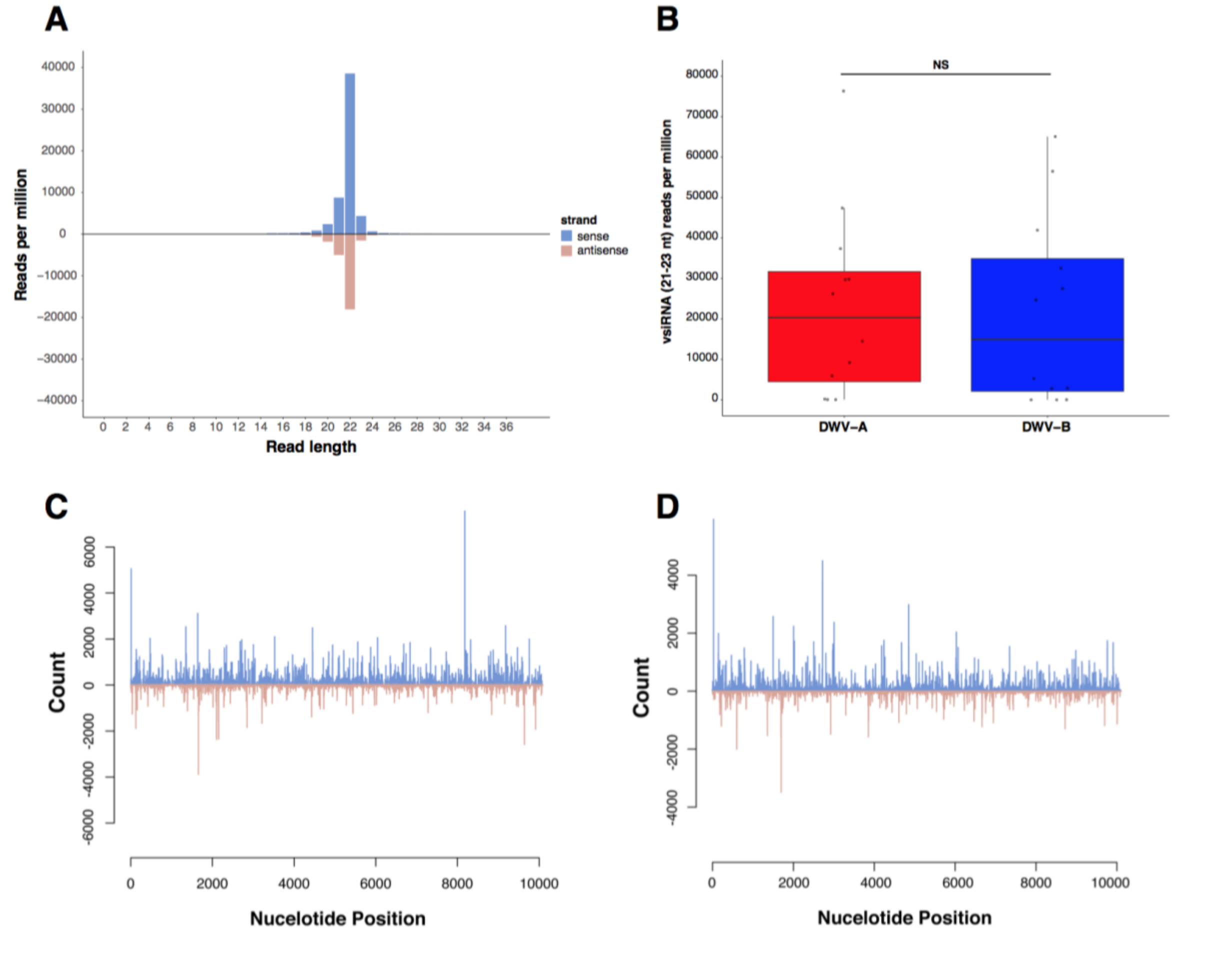


**Figure S5.** (A) Size distribution small RNA read lengths mapping to DWV. (B) Number of normalised vsiRNA (21-23 nt) reads mapping to DWV in pupae injected with DWV-A or DWV-B did not differ between the two genotypes ($\chi^2$= 0.27, df = 1, $p$ = 0.6033). Distribution of vsiRNA (21-23 nt) reads mapping to the (C) DWV-A and (D) DWV-B genomes. Data from one representative sample shown in figures 5A, 5C and 5D.

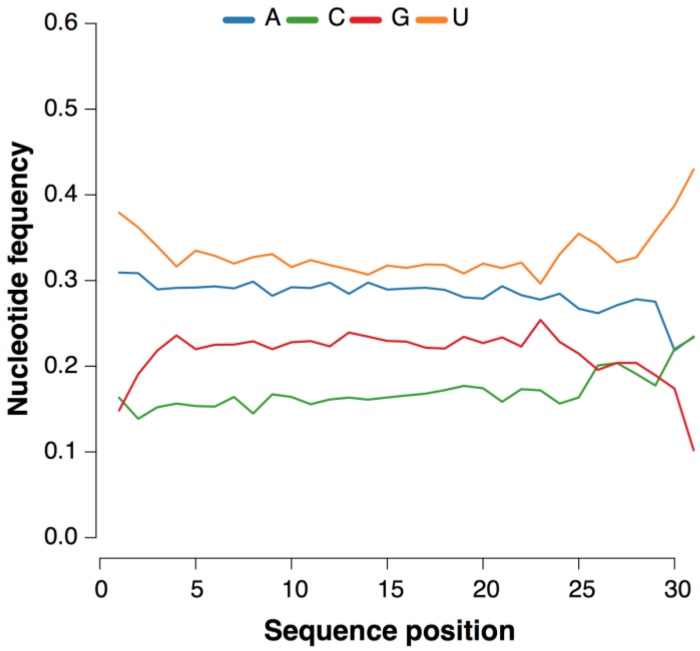


**Figure S6.** Nucleotide frequency of small 26-31 nt length RNA reads. We observed elevated frequencies of uridine at position 1, but did not observe enrichment of adenine at position 10. Thus, no evidence of a 'ping-pong' signature associated with piRNA activity was detected.

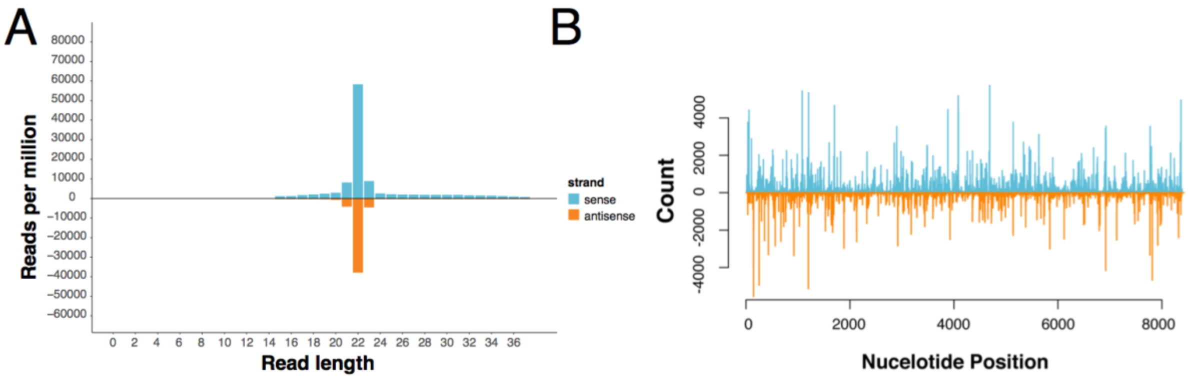


**Figure S7.** A) Size distribution small RNA read lengths mapping to BQCV. B) Distribution of vsiRNA (21-23 nt) reads mapping to the BQCV genome. Data from one representative sample shown.

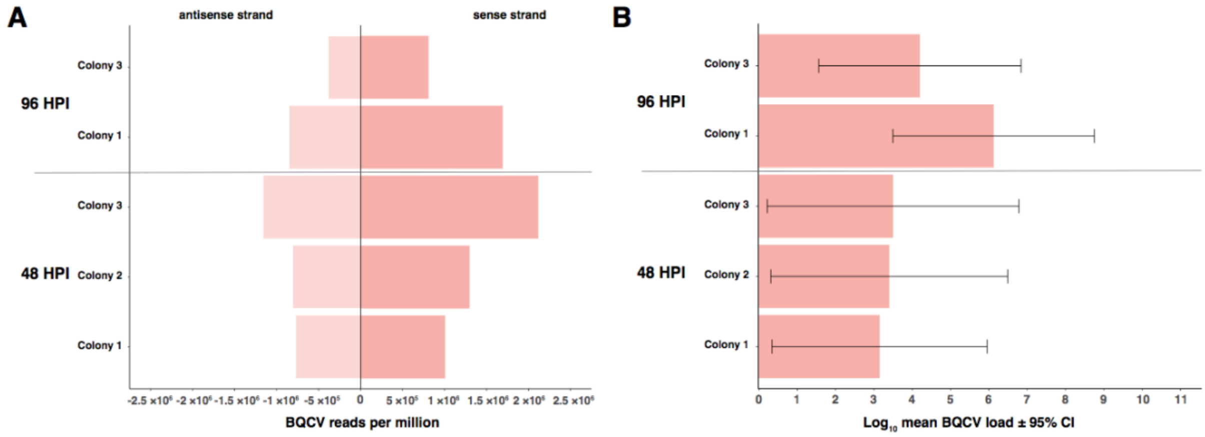


**Figure S8.** (A) Normalised siRNA (21-23 nt) reads mapping to BQCV in four pooled DWV-A injected pupae at 48 and 96 HPI, per colony. No reads mapping to BQCV were detected in the pooled sample from colony 2 at 96 HPI. (B) The mean BQCV loads ± 95% CI of the same four pupae. The large confidence intervals indicate the high variability in BQCV loads between the four pupae. Viral loads are relative to housekeeping gene *Actin* and were previously published in Norton et al. (2020), but presented here to illustrate the relationship between viral accumulation and siRNA response to BQCV. We did not detect a correlation between BQCV load and small RNA response ($r_s = 0.2$; $P = 0.7471$).
